# Comparative Annotation Toolkit (CAT) - simultaneous clade and personal genome annotation

**DOI:** 10.1101/231118

**Authors:** Ian T. Fiddes, Joel Armstrong, Mark Diekhans, Stefanie Nachtweide, Zev N. Kronenberg, Jason G. Underwood, David Gordon, Dent Earl, Thomas Keane, Evan E. Eichler, David Haussler, Mario Stanke, Benedict Paten

## Abstract

The recent introductions of low-cost, long-read, and read-cloud sequencing technologies coupled with intense efforts to develop efficient algorithms have made affordable, high-quality *de novo* sequence assembly a realistic proposition. The result is an explosion of new, ultra-contiguous genome assemblies. To compare these genomes we need robust methods for genome annotation. We describe the fully open source Comparative Annotation Toolkit (CAT), which provides a flexible way to simultaneously annotate entire clades and identify orthology relationships. We show that CAT can be used to improve annotations on the rat genome, annotate the great apes, annotate a diverse set of mammals, and annotate personal, diploid human genomes. We demonstrate the resulting discovery of novel genes, isoforms and structural variants, even in genomes as well studied as rat and the great apes, and how these annotations improve cross-species RNA expression experiments.

## Introduction

Short-read sequencing prices continue to drop and new technologies are being combined to produce assemblies of quality comparable to those previously created through intensive manual curation^1–5^ (Kronenberg et al., submitted). These advances have allowed researchers to perform clade genomics, producing assemblies for multiple members of a species or clade^6, 7^, and are required for the ambitious goals of projects such as Genome 10K^8^, which aim to produce thousands of assemblies of diverse organisms. In addition, efforts are growing to produce *de novo* assemblies of individual humans to evaluate the human health implications of structural variation and variation within regions not currently accessible with reference assisted approaches^9–11^.

These advances in genome assembly require subsequent advances in genome comparison. Central to this comparison is annotation. The challenge of finding functional elements in genome assemblies has been considered for at least the past 20 years^12^. This problem is traditionally approached by *ab initio* prediction (using statistical models of sequence composition)^13^ and sequence alignment of known mRNAs or proteins^14^. The former has limited accuracy while the latter is limited by the existence of useful sequence information. Annotation pipelines such as MAKER^15^, RefSeq^16^ and AUGUSTUS^17^ make use of both approaches. See^18^ for a recent review of genome annotation methods.

A huge amount of effort has gone into the annotation of model organisms, in particular human and mouse. For the past five years, the GENCODE Consortium^19^ has used a wide range of sequencing and phylogenetic information to manually build and curate comprehensive annotation sets, with over 43,281 and 60,297 open-reading frames in mouse and human, respectively. The GENCODE databases give a glimpse into the diversity of alternative isoforms and noncoding transcripts present in vertebrate genomes. Similarly, efforts in other model organisms, such as zebrafish^20^, *C. elegans*^21^, *A. thaliana*^22^ and many others, have produced high-quality annotation sets for their respective assemblies.

As we enter a third era of genome assembly, consideration should be given to scaling annotation. Here, we present a method and toolkit to make use of multiple genome alignments produced by Progressive Cactus^23^ and existing high-quality annotation sets to simultaneously project well-curated annotations onto lesser studied genomes. In contrast to most earlier alignment methods^24–26^, Progressive Cactus alignments are not reference based, include duplications, and are thus suitable for the annotation of many-to-many orthology relationships. We show how the output of this projected annotation set can be cleaned up and filtered through special application of AUGUSTUS^27^ and how novel information can be introduced by combining the projected annotation set with predictions produced by Comparative Augustus^28^. These predictions can be further supplemented and validated by incorporating long-range RNA-sequencing (RNA-seq) data, such as those generated by the Iso-Seq protocol^29^. We provide a fully featured annotation pipeline, the Comparative Annotation Toolkit (CAT), that can perform this annotation process reproducibly on any combination of a local computer, a compute cluster, or on the cloud. We show that this pipeline can be applied to a wide range of genetic distances, from distant members of the same clade to individualized assemblies of the same species.

## Results

### Comparative Annotation Toolkit

CAT provides a software toolkit designed to perform end-to-end annotation; Figure 1 gives an overview. The only required inputs are a hierarchical alignment format (HAL)^30^ multiple genome alignment as produced by Progressive Cactus and a GFF3 format annotation file for the previously annotated genome(s). CAT can take as optional input a set of aligned RNA-seq or Iso-Seq BAM format files, as well as protein FASTA files, which are used to construct hints for AUGUSTUS.

**Figure 1.**
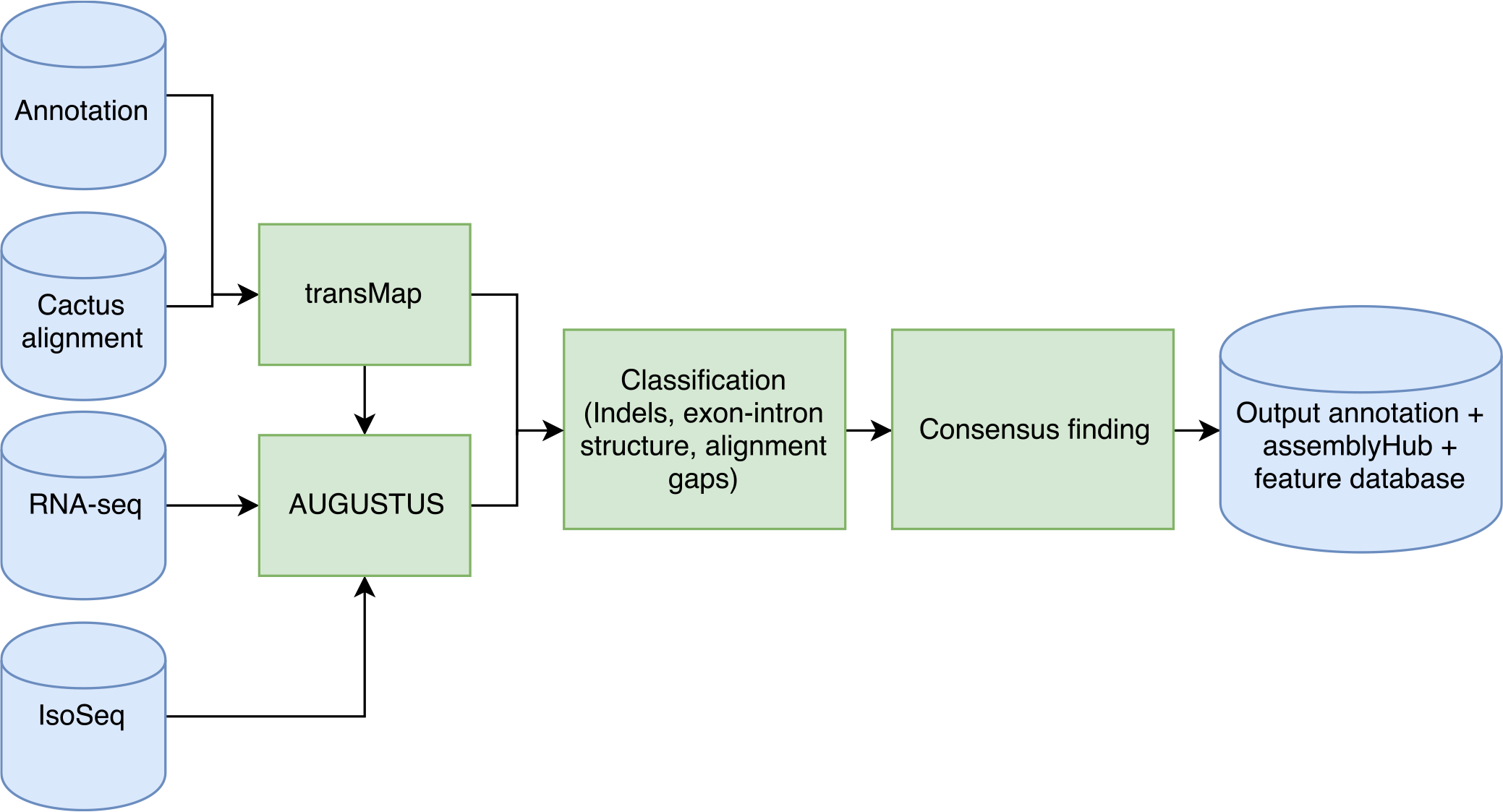
CAT pipeline schematic. The CAT pipeline takes as input a HAL alignment file, an existing annotation set and aligned RNA-seq reads. CAT uses the Cactus alignment to project annotations to other genomes using transMap^27^. These transcript projections are then filtered and paralog resolved. Optionally, AUGUSTUS can be run into up to four parameterizations. All transcripts are classified for extrinsic support and structure and a ‘chooser’ algorithm picks the best representative for each input transcript, incorporating *ab-initio* transcripts when they provide novel supported information. The final consensus gene set as well as associated feature tracks are used to create a assembly hub ready to be loaded by the UCSC Genome Browser. See Supplementary Figure S1 and Supplementary Note 1 for more detail.

TransMap^27, 31^ is used to project existing annotations between genomes using the Progressive Cactus alignment. TransMap projections are filtered based on a user-tunable flag for minimum coverage, and then the single highest scoring alignment is chosen. If this results in transcripts for a given gene mapping to multiple loci, these are resolved to one locus based on the highest average score of a locus, rescuing lower scoring alignments.

Based on input parameters, CAT will run AUGUSTUS in up to four distinct parameterizations, two of which rely on transMap projections (AugustusTMR) and two that perform *ab initio* predictions (AugustusCGP and AugustusPB) using extrinsic information to guide prediction. AugustusCGP performs simultaneous comparative prediction^28^ on all aligned genomes, while AugustusPB uses long read RNA-seq to discover novel isoforms. The output of these modes of AUGUSTUS are evaluated alongside the original transMap projections using a combination of classifiers as well as the output from homGeneMapping^13^, which uses the Cactus alignments to project features such as annotations and RNA-seq support between the input genomes. AugustusCGP and AugustusPB transcript predictions are assigned to transMap genes based on genomic and exonic overlap. If they overlap projections that were filtered out in the paralog resolution process, then they are flagged as putatively paralogous, while if they do not overlap any transMap projections they are flagged as putatively novel. A consensus-finding algorithm combines these gene sets.

The consensus finding algorithm combines all sources of transcript evidence into an annotation set. On a per-gene basis, it evaluates the transMap transcripts for passing user tunable flags for RNA-seq and annotation support. It then considers the inclusion of *ab initio* transcripts based on their assignment to this locus and their contribution of novel splice junctions supported by RNA-seq or IsoSeq. Finally, it evaluates *ab inito* transcripts not assigned to a gene as novel loci if they are supported by RNA-seq or IsoSeq as defined by user-tunable flags. For a more detailed description of CAT see Supplementary Note 1.

### Annotation of great apes

The previous generation of great ape assemblies (panTro4, ponAbe2 and gorGor4) as well as the new SMRT (PacBio) great ape assemblies^5^ (Kronenberg et al., submitted) were annotated by CAT, using GRCh38 and GENCODE V27 as the reference. On average, CAT identified 141,477 more transcripts and 25,090 more genes in the new SMRT assemblies of the great apes compared to the Ensembl V91 annotation of the previous generation of great ape assemblies. Relative to the existing human annotation, the CAT annotations represent an average of 95.0% of GENCODE gene models and 94.3% of GENCODE isoforms in the SMRT great ape assemblies. This increase in isoform representation is mostly due to the large number of isoforms annotated by GENCODE and reproduced in these genomes, while the increase in gene content is due to the mapping over of non-coding genes poorly represented in the Ensembl annotation. Comparing the CAT annotations of SMRT great apes and older assemblies, we see an average increase of 610 genes (1.9%) and 3,743 isoforms (1.0%) (Supplemental Figure S2) in the SMRT assemblies; given this relatively small increase, most of the observed increase in genes and isoforms in the CAT annotations relative to the Ensembl annotations are therefore a result of the CAT annotation process rather than the updated assemblies.

Conversely to the overall increases in genes and isoforms, CAT identifies on average 3,553 fewer protein-coding genes than Ensembl. However, this brings the total number of coding genes more closely in line with the GENCODE annotation of human, as Ensembl has an average of 2,081 more protein-coding genes in great apes than GENCODE has for human (Supplemental Figure S2).

To evaluate these annotations in a non-species-biased fashion, consensus isoform sequences created from Iso-Seq reads for each species were compared to their respective species annotations. As a baseline comparison, equivalent human data was compared to the high-quality human GENCODE V27 annotation. The CAT annotation of both the SMRT and older great ape assemblies (which used the raw Iso-Seq reads during the annotation process) and the Ensembl annotation of the older assemblies were compared. We calculated the rate of isoform concordance, that is the fraction of consensus Iso-Seq sequences that match either exactly or fuzzily an annotated isoform (Figure 2A; methods). Fuzzy matching allows for the intron boundaries to shift slightly in a isoform. For the SMRT chimpanzee (74.0%/82.1% exact/fuzzy matching) and orangutan (71.4%/80.4%) genome assemblies the isoform concordance rates were comparable to the rate for human (74.6%/82.1%). The gorilla GSMRT3.2 assembly showed lower concordance (67.6%/76.9%), likely due to the higher indel error rate in that assembly (Supplemental Figure S3). In contrast, the isoform concordance rate for the older assemblies was lower (on average 60.0%/69.6%), mostly reflecting exons in gaps and mis-joins, and was lower still for the existing Ensembl annotations (on average 47.9%/57.6%).

**Figure 2.**
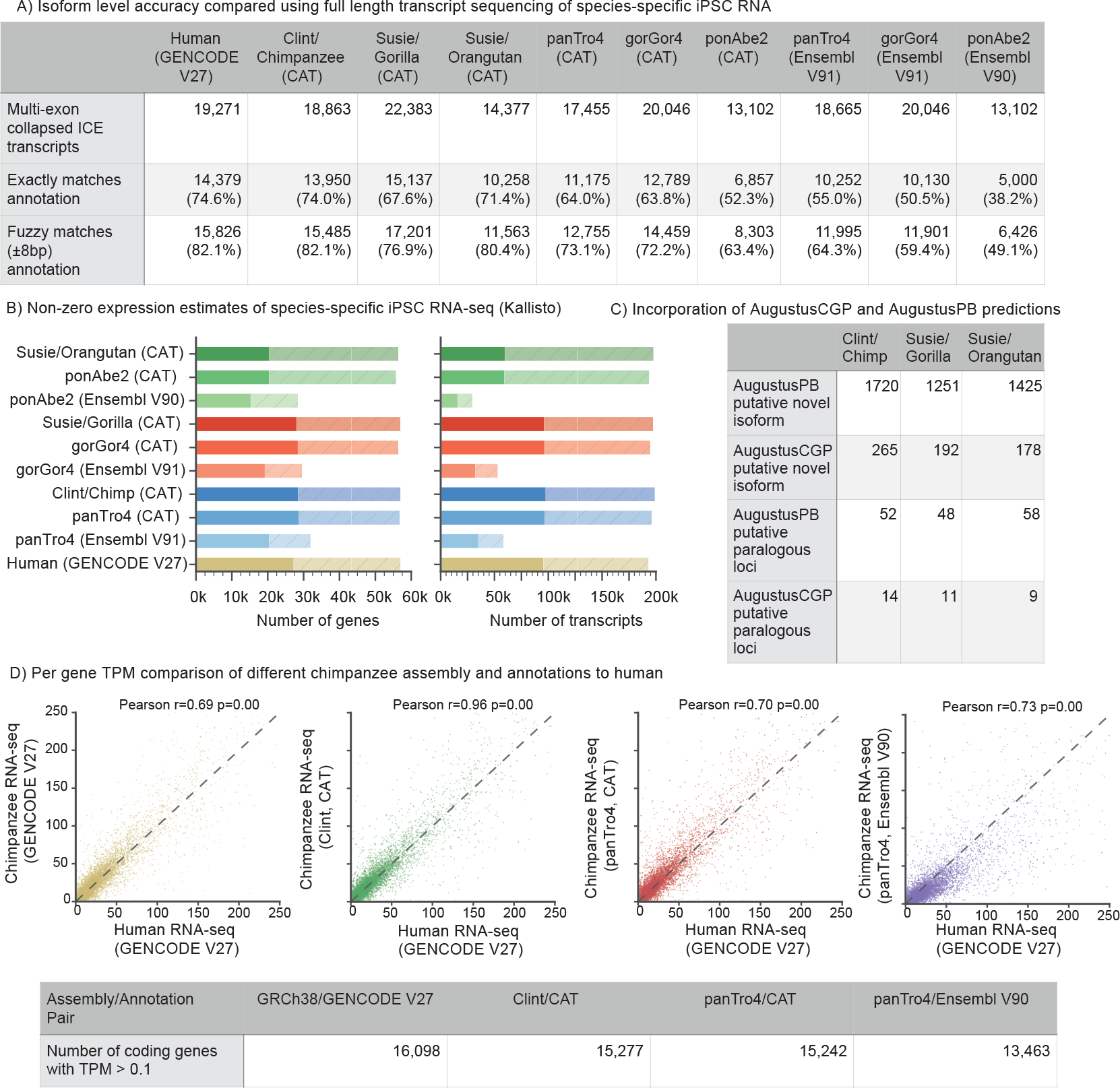
Primate annotation. A) Validating CAT annotations using Iso-Seq data. As a baseline comparison, Iso-Seq data from human iPSCs were compared to the GENCODE V27 annotation. Iso-Seq data from chimpanzee, gorilla and orangutan iPSC lines were compared to respective species-specific annotations. The Iso-Seq data were clustered with ICE and collapsed using ToFu^29^. CAT annotation of PacBio great apes showed similar isoform concordance to human, and improvement over the older assemblies. B) Kallisto^32^ was used to quantify liver Illumina RNA-seq from each species on both the gene and transcript level on the existing and new great ape assemblies. Solid bars are transcripts or genes with transcripts per million (TPM) *>*0.1, while shaded hatched bars are the remainder of the annotation sets. CAT annotation of great apes shows nearly the same number of expressed genes and isoforms as the GENCODE reference on human with the exception of orangutan. C) The number of novel isoforms and paralogous genes with Iso-Seq support discovered by analysis of AugustusPB and AugustusCGP predictions for each species. D) Kallisto protein-coding gene-level expression for chimpanzee iPSC RNA-seq is compared to human across all of the chimpanzee annotation and assembly combinations as well as when mapped directly to human. In all cases the x-axis is the TPM of human iPSC data mapped to human. The highest correlation (Pearson r=0.96) is seen when comparing Clint annotated with CAT to GRCh38 annotated with GENCODE V27.

To assess the utility of CAT annotations for short-read analysis of RNA expression, species-specific induced pluripontent stem cell (iPSC) Illumina RNA-seq data were quantified (Figure 2B). Comparing the annotations of the older assemblies, CAT identified an average of 9,518 more genes and 54,107 more transcripts with measurable expression compared to Ensembl.

We might expect the per-gene abundance estimates of the majority of genes in matched cell types to agree between species, particularly for closely related species. It is reasonable to therefore prefer *a priori* an annotation of the great apes that produces expression estimates that agree with expression estimates from the matched human data using the GENCODE annotation. Doing these comparisons, we find better correlations on average using the CAT annotation of the older assemblies (avg. Pearson r=0.63; Figure 2D, Supplemental Figure S4) than the Ensembl annotations of the older assemblies (avg. Pearson r=0.44). However, we find by far the highest correlation when CAT annotates the SMRT primate assemblies (avg. Pearson r=0.90). This reflects the increased representation in the updated assemblies of transcript sequence, especially 3’ UTRs that are important for quantifying polyA primed RNA-seq (Kronenberg et al., submitted). Notably, we find that the correlations between the CAT annotations of the SMRT assemblies and the matched human data are higher than when mapping the species-specific data back to the human GENCODE annotations and comparing to the human data (Figure 2D, Supplemental Figure S4), demonstrating the benefit of having species-specific annotations within closely related species that have clear cross-species orthology relationships. Analysis at the isoform level showed the same patterns (Supplemental Figure S5), albeit with slightly weaker correlations.

Predictions performed by AugustusCGP and AugustusPB were incorporated into the gene sets based on the presence of splice junctions supported by RNA-seq or Iso-Seq and not present in the transMap/AugustusTMR-derived annotations (Figure 2C). An average of 1,677 novel isoforms and 64 novel loci were found across the assemblies with at least one Iso-Seq read supporting the prediction.

CAT provides new metrics for diagnosing assembly quality. In the process of annotating the great ape genomes, we noticed that assemblies that had undergone Quiver and Pilon^33^ correction still exhibited a systematic bias towards coding deletions. These were identified to be related to heterozygosity in the input dataset and a variant-calling-based correction method (Kronenberg et al., submitted) was developed to resolve these issues, dramatically lowering the coding indel rate and reducing systematic bias (Supplemental Figure S3). CAT can also measure gene assembly contiguity by reporting the number of genes whose transcripts end up split across multiple contigs, or on disjoint intervals in the same contig. Comparison of split gene metrics between the old and new primate assemblies shows 504 fewer split genes in chimpanzee, 560 fewer in gorilla and 1,858 fewer in orangutan (Supplemental Figure S6).

### Annotation of personal human diploid assemblies

High-quality *de novo* assembly of a human genome is increasingly feasible; both Pacific Biosciences^34–36^ (Falcon) and 10x Genomics^37^ (Supernova) provide tools to construct phased, diploid assemblies. Annotating diploid assemblies provides a window into haplotype-specific structural variation that may affect gene expression. To evaluate the ability of CAT to provide this analysis, Progressive Cactus alignments were generated between hg38 and the two haploid cell line assemblies, CHM1 (GCA 001297185.1) and CHM13 (GCA 000983455.2) as well as the 10x Genomics diploid assemblies of four individuals (NA12878, NA24385, HG00512 and NA19240).

An average of 98.5% of genes present in GENCODE V27 were identified in CHM1/CHM13, while an average of 97.3% of genes were identified in the 10x Supernova assemblies. (Figure 3A). After filtering, an average of 552 genes in the PacBio assemblies and 461 genes in the 10x assemblies had frame-shifting indels (Figure 3B). Compared to ExAC, which found between 75 and 125 putative truncating events per individual^38^, this result suggests indel errors in the assemblies are producing false positives. All assemblies exhibit systematic overrepresentation of deletions, including the Pacbio assemblies despite coming from haploid cell lines (Figure 3B). Split gene analysis found the CHM1 assembly to be the most gene contiguous, with only 39 genes split across multiple contigs, and the PacBio assemblies overall more contiguous (Figure 3C). Gene contiguity is measured by looking at genes with multiple alignments post paralog resolution that start and end nearby in transcript coordinates.

**Figure 3.**
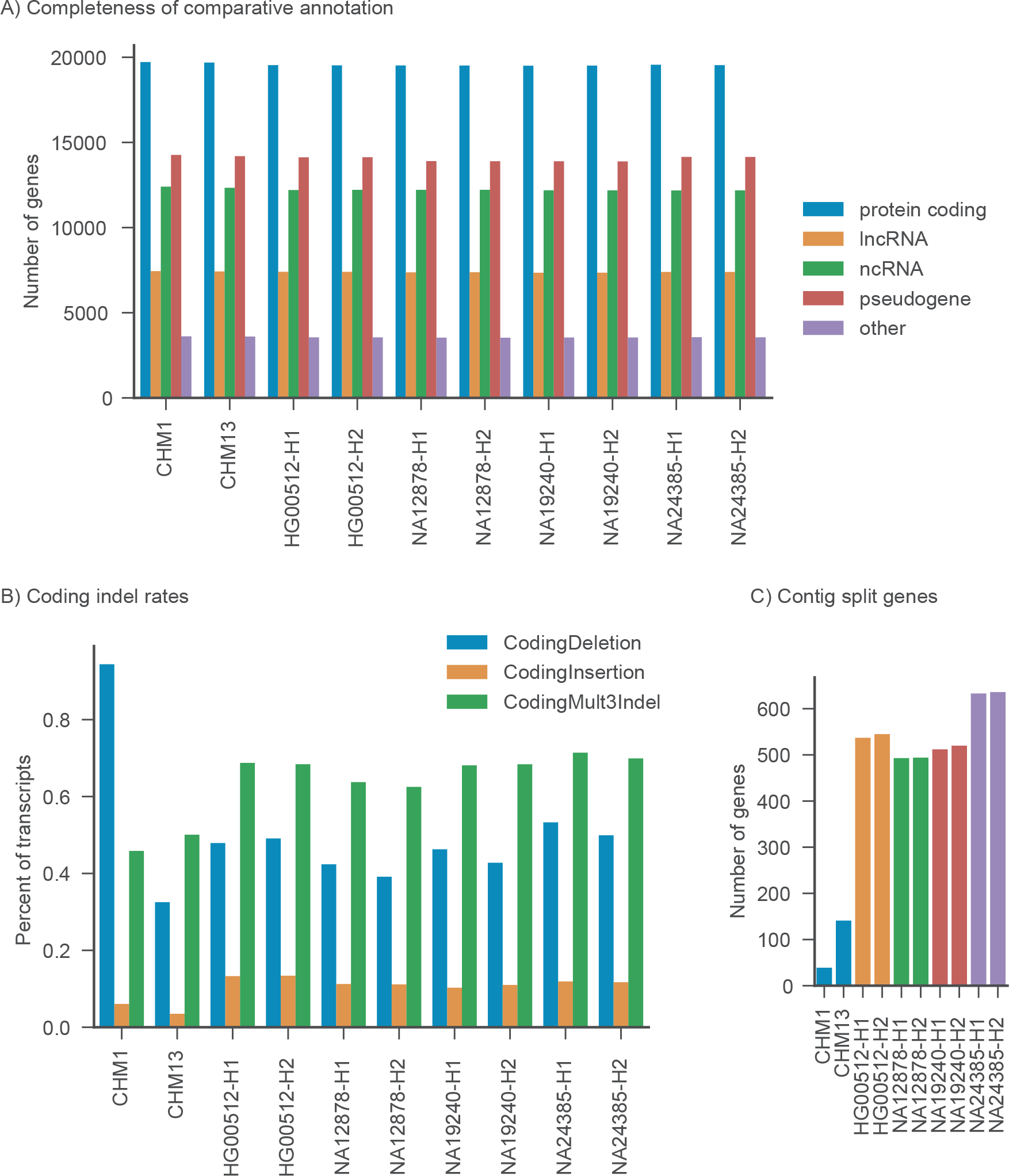
Pseudo-diploid human annotation metrics. A) The number and fraction of genes comparatively annotated from GENCODE V27 in each assembly. GENCODE biotypes are simplified into protein coding, lncRNA, ncRNA, pseudogene and other. Other includes processed transcripts, nonsense-mediated decay, and immune-related genes. B) Frame-shifting insertions, deletions and multiple of 3 indels that do not shift frame are reported for each assembly. Consistent with the great ape genomes, there is a systematic over-representation of coding deletions in Falcon assemblies, despite these assemblies coming from haploid cell lines. 10x Supernova assemblies also exhibit similar properties. C) Split gene analysis reports how often paralog-resolved transcript projections end up on different contigs, which can measure assembly gene-level contiguity. PacBio assemblies, especially CHM1, are the most contiguous.

Manual analysis of genes with different transMap coverage in CHM13 relative to CHM1 led to the discovery of the example region in Figure 4A. This deletion removes most of the exons of *TRIB3*, a pseudokinase associated with type 2 diabetes^39^ (Supplemental Figure S7). Similar analysis in the diploid assembly of NA12878 led to the discovery of a tandem duplication involving an exon of TAOK3 in one haplotype.

**Figure 4.**
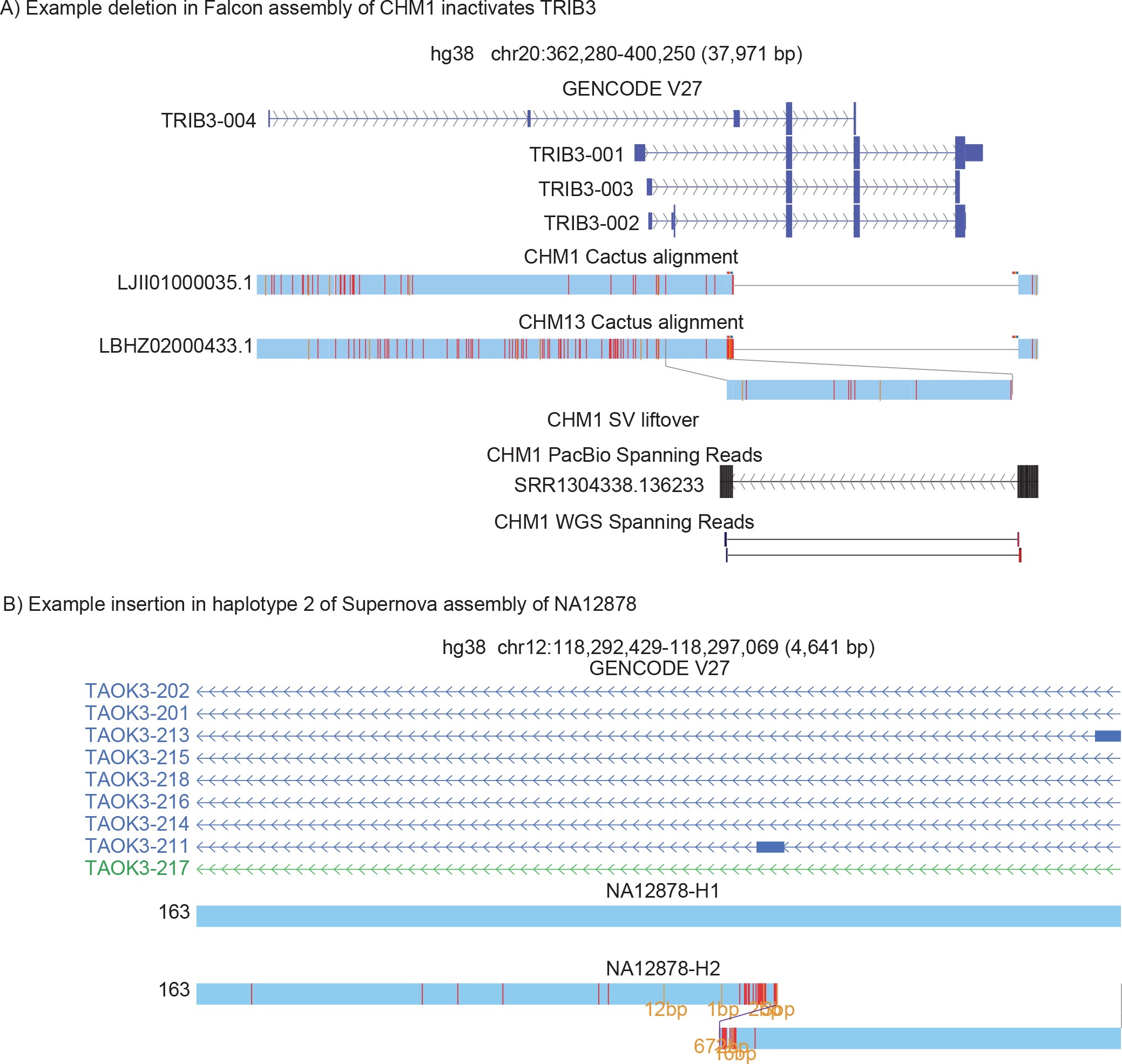
Pseudo-diploid human annotation examples. A) UCSC Assembly Hub^40^ showing TRIB3 deletion in CHM1. Analysis of genes found in one genome and not the other led to the discovery of a novel structural variant specific to CHM1, which disables the gene TRIB3. Spanning reads were found in both PacBio and Illumina whole-genome sequencing that validate the deletion. B) An example insertion near an exon of *TAOK3* seen in one haplotype of NA12878. It was not possible to determine if this insertion effected transcription of this gene.

### Reannotating the rat genome

We tested CAT’s ability to reannotate the rat genome using information from the mouse genome. These genomes differ by approximately 0.18 substitutions/site, much more, for example, than the 0.04 substitutions/site separating the human and orangutan genomes^41^.

CAT was run on a Cactus alignment between mouse (mm10) and rat (rn6) using rabbit (oryCun2), Egyptian jerboa (jacJac1) and human (hg38) as outgroups. To provide hints to AUGUSTUS, RNA-seq data were obtained from SRA^42–44^ (Supplemental Table S2). For comparison we used existing Ensembl and RefSeq rat annotations and ran the MAKER2 pipeline^15^ to generate an annotation set. MAKER was provided both a Trinity^45^ *de novo* assembly of the input RNA-seq data provided to CAT (MAKER does not process raw RNA-seq) as well as the mouse protein sequences from GENCODE VM11, together providing a comparable input set to what CAT had.

CAT comparatively annotated 78.1% of genes and 91.9% of protein-coding genes present in GENCODE VM11 on rn6 (Supplemental Figure S8), representing an increase of 14,675 genes and 74,308 transcripts over Ensembl V90, 5,104 genes and 32,157 transcripts over RefSeq and 14,541 genes and 81,022 transcripts over Maker. 13,651 loci were identified with no overlap to any other annotation set (Supplemental Figure S9).

We compared CDS exon and CDS intron predictions between annotation sets (Table 1A, Supplemental Figure S10). We measured precision and recall of coding intron and exon intervals based on comparing the CAT annotation to EnsemblV90, where precision is the proportion of CAT exons/introns that exactly match Ensembl, and recall is the proportion of CAT exons/introns that exactly match Ensembl over the number of exons/introns CAT annotated. Ensembl, RefSeq and CAT CDS exon annotations were all comparably similar (between 0.648 and 0.659 Jaccard similarity), while for CDS introns CAT and RefSeq displayed the highest Jaccard similarity (0.841). In all comparisons MAKER was the outlier (Table 1B) with the lowest similarity to the other sets.

**Table 1.**
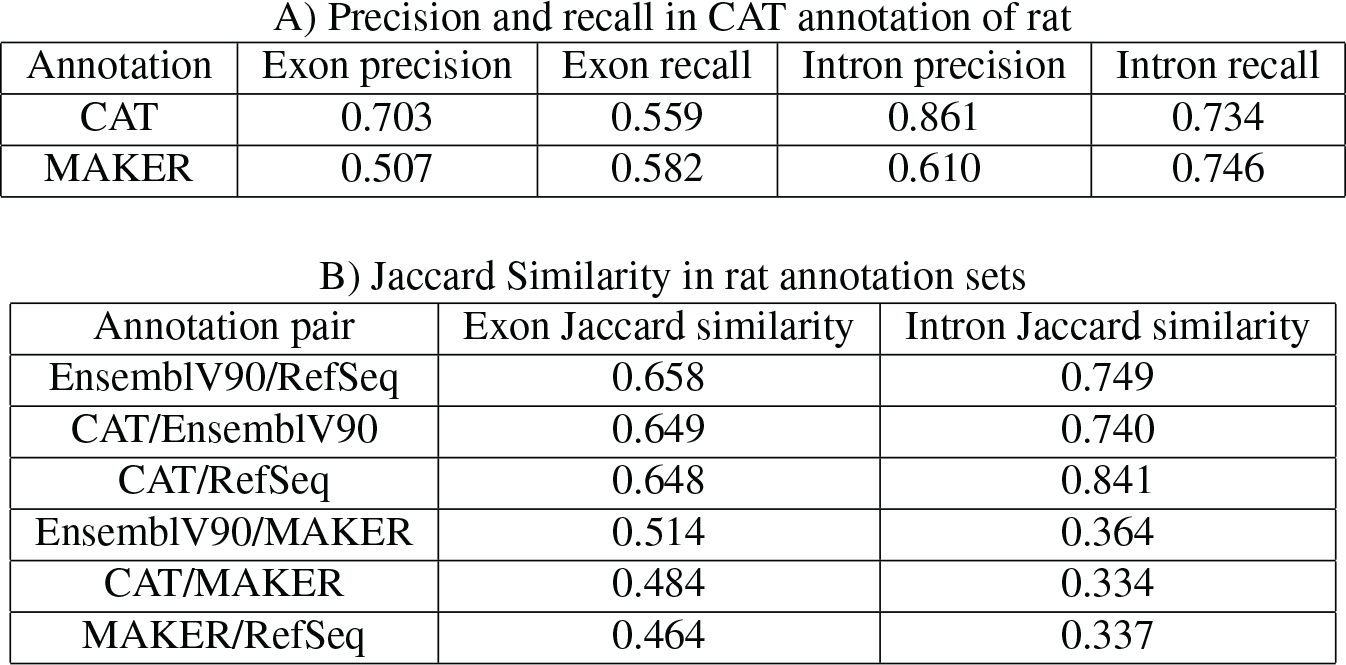
Rat annotation similarity metrics. A) Precision is the number of coding exons or coding introns that exactly match divided by the number of exons or introns in the Ensembl annotation, while recall is the number that exactly match divided by the number of exons or introns in the CAT or, respectively, MAKER annotation. B) Jaccard similarity of CDS introns and exons between rat annotation sets shows high similarity between CAT and existing Ensembl and RefSeq annotations, comparable to the similarity between Ensembl and RefSeq themselves.

**A) Precision and recall in CAT annotation of rat**
| Annotation | Exon precision | Exon recall | Intron precision | Intron recall |
| --- | --- | --- | --- | --- |
| CAT | 0.703 | 0.559 | 0.861 | 0.734 |
| MAKER | 0.507 | 0.582 | 0.610 | 0.746 |

**Table 1.** Rat annotation similarity metrics
| Annotation pair | Exon Jaccard similarity | Intron Jaccard similarity |
| --- | --- | --- |
| EnsemblV90/RefSeq | 0.658 | 0.749 |
| CAT/EnsemblV90 | 0.649 | 0.740 |
| CAT/RefSeq | 0.648 | 0.841 |
| EnsemblV90/MAKER | 0.514 | 0.364 |
| CAT/MAKER | 0.484 | 0.334 |
| MAKER/RefSeq | 0.464 | 0.337 |

The input RNA-seq dataset was used for isoform quantification against the CAT, MAKER, Ensembl and RefSeq transcriptomes (Figure 5A). CAT identified 1,881 protein-coding genes and 1,011 lncRNAs with measurable expression not present in either Ensembl or RefSeq. CAT also identified 27,712 expressed coding splice junctions not in the union of RefSeq and Ensembl, for a total of 21,267 novel expressed isoforms. 5,526 of the 13,651 loci unique to CAT had measurable expression.

**Figure 5.**
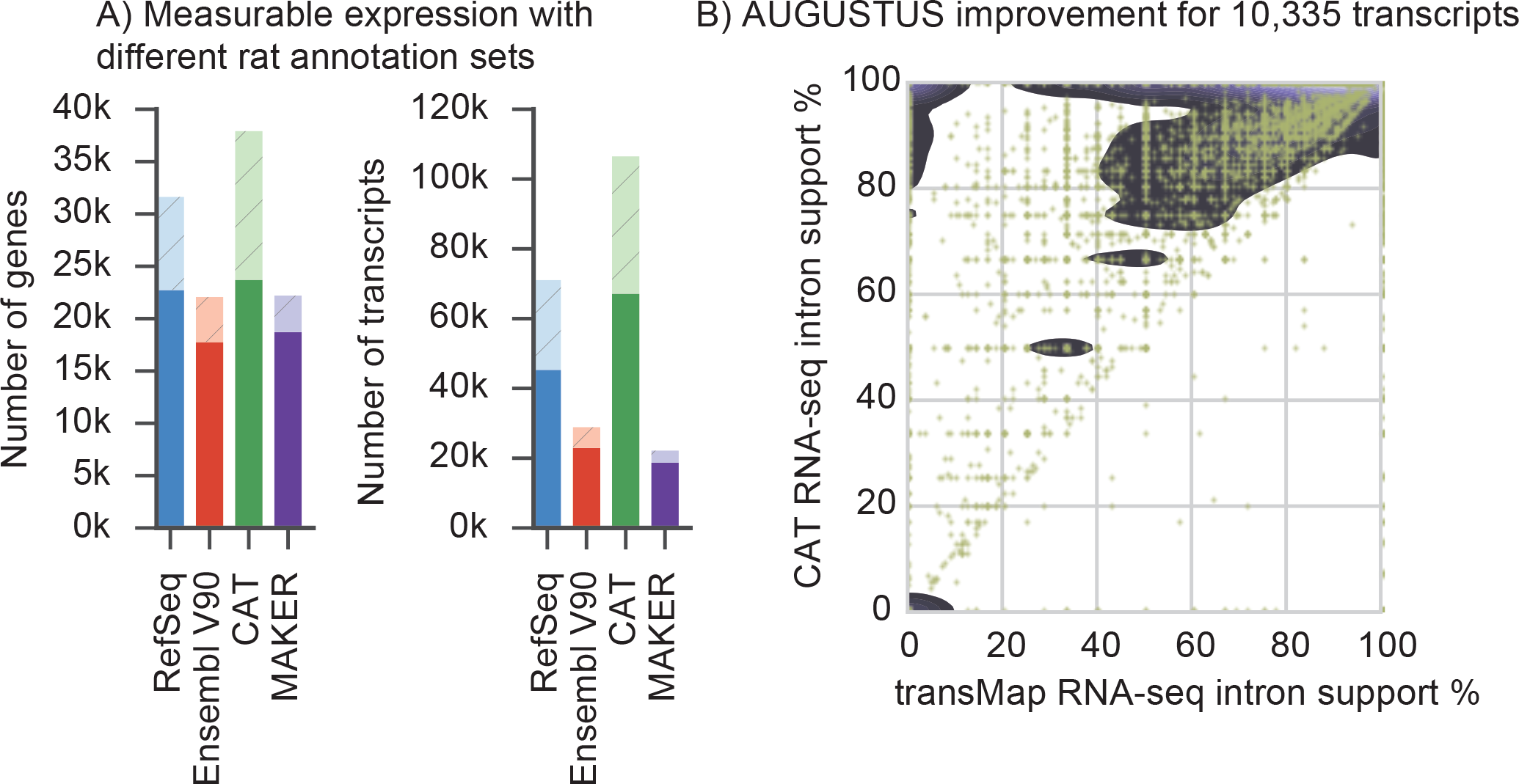
Validation of CAT annotation using rat. A) Each transcript set was used to construct a Kallisto^32^ index, and then all of the input RNA-seq for annotation were quantified. Solid bars are genes or transcripts with non-zero expression (TPM*>*0.1) estimates, while light hatched bars are the remainder of the annotation set. CAT provides an annotation set with slightly more detectable genes than other annotation methods, and far more detectable isoforms. B) AugustusTMR provides a mechanism to clean up transcript projections and shift splice sites, fixing alignment errors as well as real evolutionary changes. Most of the 10,335 AugustusTMR transcripts chosen in consensus finding show an improvement in RNA-seq support, which is one of the features used in consensus finding.

AugustusTMR, which uses transMap and RNA-seq, provides CAT with a way to improve transcript predictions projected between species. Comparing the 10,335 multi-exon protein-coding transcripts in which the AugustusTMR prediction differed from the input transMap projection, we see considerable overall improvement in resulting RNA-seq support of predicted splice boundaries in the AugustusTMR transcripts (Figure 5B).

### Annotation of a diverse set of mammals

Finally, to test CAT’s ability to annotate across a substantial and diverse range of genomes, 13 mammalian genomes were comparatively annotated using the mouse (mm10) GENCODE VM15 as the reference transcript set (Figure 6A). Species-specific RNA-seq was used for every genome (Supplemental Table S2). To assess the completeness of these annotation sets, 4,104 benchmarking universal single-copy orthologs (BUSCO) were used^46^, which by design should be nearly uniformly present in each of these genomes. On average, only 108 BUSCO genes (2.63%) were not annotated by CAT in each genome (Supplementary Table S1).

**Figure 6.**
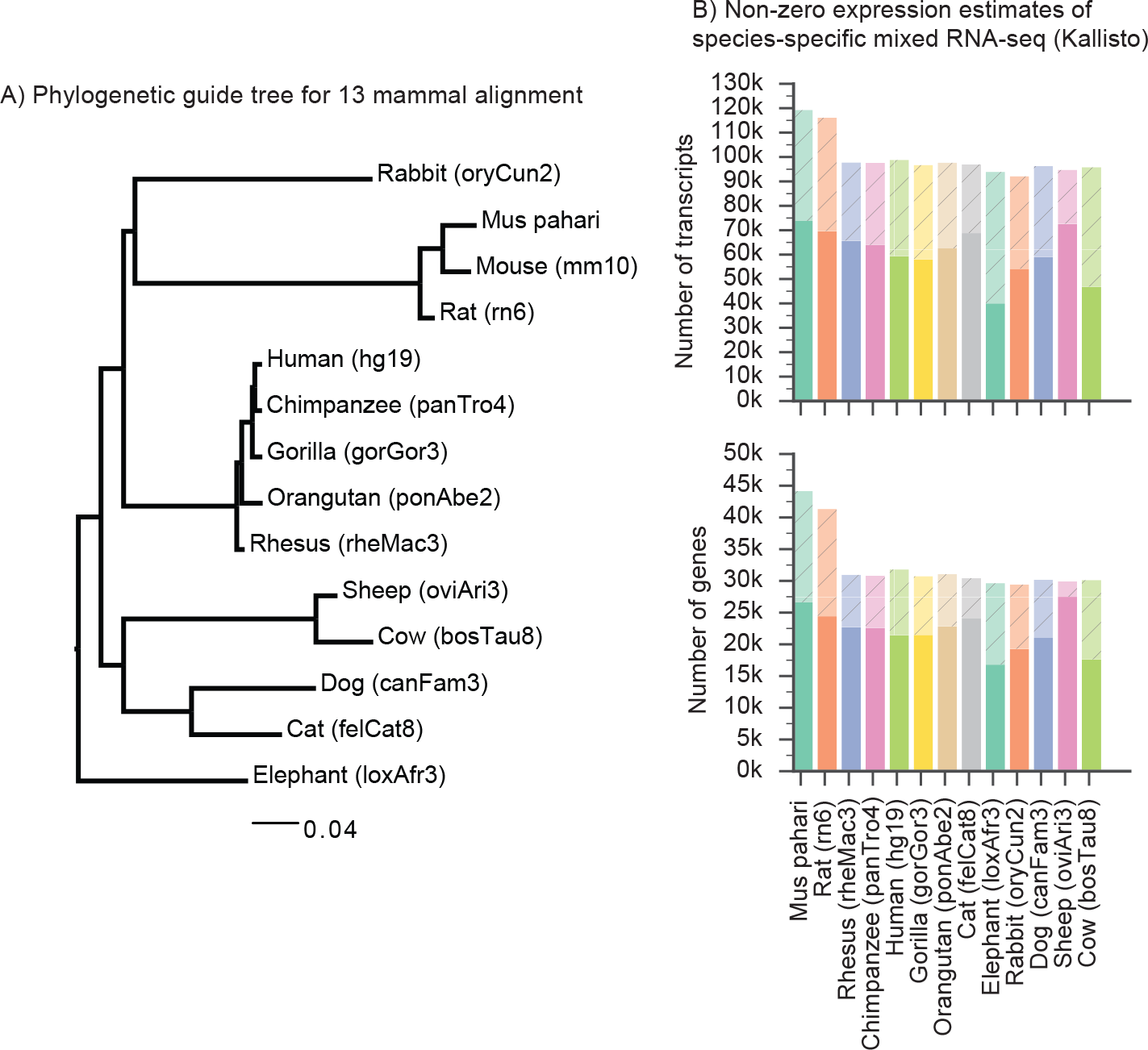
13-way annotation. A) The phylogenetic guide tree for the 14-way mammal alignment. See the methods section for the exact Newick format tree. B) The gene annotation sets for each of the 13 mammalian genomes were quantified against the mixed input RNA-seq sets obtained from SRA. Genes or transcripts with TPM*>*0.1 are solid colors, while genes or transcripts with no measurable expression are shaded. An average of 2.8 isoforms per gene per genome had quantifiable expression, suggesting that CAT can infer isoform information across long branch lengths.

To estimate the usefulness of these annotation sets, the input RNA-seq datasets were used to quantify expression of the annotation sets (Figure 6B). The main factor in measurable expression is the variety of the input RNA-seq datasets, as exemplified by the ability to measure expression of 88.9% of genes annotated in the sheep genome.

To assess the CAT translation of annotations over large phylogenetic distances, as well as provide a baseline validation of CAT performance, the annotation of human hg19 (GRCh37) produced in the representative mammalian genome annotation was compared to the current human GENCODE annotation set for that assembly (GENCODE V27lift37). Of the 19,233 ICE isoforms detected when running ToFu^29^ against hg19, 12,911 (67.2%) fuzzy matched a CAT isoform compared to 15,920 (82.8%) of the human GENCODE annotations. Precision and recall analysis shows results similar to the rat annotation, with better matches in introns. 91.2% of CAT introns and 75.2% of CAT protein coding isoforms match GENCODE (Table 2).

**Table 2.** Precision and recall of CAT annotation of hg19 using mouse isoforms Precision and recall are measured by looking at exact matches of coding introns, exons and isoforms. Isoforms are compared on a coding intron chain level. Precision and recall are defined in the same way as in Table 1.

| Exon precision | Exon recall | Intron precision | Intron recall | Isoform precision | Isoform recall |
| --- | --- | --- | --- | --- | --- |
| 0.532 | 0.688 | 0.777 | 0.912 | 0.408 | 0.752 |

## Discussion

Gene annotation is a longstanding and critical task in genome informatics that must now be scaled to handle the rapidly increasing number of available genomes. At the time of writing there were 570 vertebrate genomes available from NCBI, but only 100 (17.5%) and 237 (41.6%) had Ensembl and RefSeq annotations, respectively.

We introduce CAT to help meet this need, building around a number of key innovations. Firstly, CAT utilizes the reference-free, duplication-aware multiple genome alignments we have developed. This allows CAT to annotate multiple genomes symmetrically and simultaneously, breaking from the traditional pattern of annotating each new genome individually, as is currently the practice for the RefSeq, Ensembl and MAKER gene-building pipelines. Not only does this solve a key scalability issue by annotating multiple genomes simultaneously and consistently, CAT is able to produce orthology mappings, naming each equivalence class of orthologs based upon an initial reference annotation, and add to this sets of newly discovered genes. This can provide valuable evolutionary insights. For example, the analysis of the rat genome shows that many of the alternative isoforms and projected transcription start sites identified by GENCODE in mouse genes are supported by expression analysis in rat (Supplemental Figure S11).

A second key innovation made by CAT is its leveraging of existing reference annotations. A huge amount of effort has been placed into the annotation of key species, such as human and mouse, employing a myriad of technologies and extensive, labor-intensive manual curation. It is very unlikely that this effort will be replicated across a significant fraction of other genomes, so instead we propose the ‘project and augment’ strategy employed by CAT to annotate related genomes. Here we show that this strategy is very clearly able to improve the annotation of great ape genomes, using the human GENCODE set as the reference, and we make the case that we can even improve the annotation of a genome as well studied as the rat.

To circumvent the reference bias of existing annotations and to discover new genes and isoforms, CAT is able to integrate multiple forms of extrinsic information, using multiple, novel parameterizations of the AUGUSTUS algorithms. This includes use of new long-read RNA data, in particular Iso-Seq data, and shortly will integrate nanopore-based long-read data^47^. Using this expression data not only allowed us to confirm expression of a substantial fraction of isoforms but allowed us to discover thousands of novel isoforms and dozens of novel genes in the great ape genomes.

With the advent of more affordable *de novo* genome assembly, there is renewed interest in the generation of *de novo* human genomes, and in general the creation of multiple *de novo* genomes for a species. This has the advantage of providing fully independent reconstruction and is particularly appropriate for sequences that are highly divergent from the reference, e.g. structural variations. However, such assemblies do not negate the need for genome comparison; Cactus can be parameterized to rapidly create sensitive whole-genome alignments of human genomes, and here we have demonstrated that CAT can be used to build upon this to produce a high-quality diploid gene annotation and ortholog mapping.

CAT works best when provided RNA-seq data, but for many species this may not be possible. From our experience, a reasonable amount (on the order of 50 million reads) of RNA-seq from tissues like brain and liver is fairly informative. Using polyA-selected libraries is recommended, as it greatly reduces false positive predictions in AugustusCGP. Iso-Seq data allowed for the discovery of thousands of novel isoforms in the great apes but may be too expensive for many projects. In clade genomics projects, we would suggest generating RNA-seq for a few of the species and then reliance on the coordinate mapping that AugustusCGP and homGeneMapping provide to evaluate support in other members of the clade.

A key barrier to the use of bioinformatics tools is their ease of use; we have focused on providing cloud agnostic distributions of the CAT software so that, despite its complexity, it can be run within a uniform computational environment by external groups.

CAT is not without limitations. In the future it would be good to use the genome alignments to not only project transcripts, but to use the evolutionary conservation signatures to predict the potential likelihood of projected annotations being coded^48^. CAT also does not yet provide means to detect new processed, unprocessed and unitary pseudogene predictions other than via projection of existing annotations. CAT’s current implementation also does not attempt to put weights on the features used for constructing a consensus gene set. Instead, it simply scores transcripts based on the sum of all features evaluated. In the future deep learning methods could be added to CAT to construct feature weights and improve consensus finding, better mimicking the labor intensive efforts of manual annotators who currently weigh such evidence.

An earlier version of CAT was used to annotate the PacBio-based assembly of the gorilla genome^5^ as well as produce the current Ensembl annotations for 16 laboratory mouse strains as part of the Mouse Genomes Project^49^ (http://www.sanger.ac.uk/science/data/mouse-genomes-project). In addition, CAT has been proposed for the Vertebrate Genomes Project (VGP), which aims as a pilot project to assemble and annotate one member of every order of vertebrate species. CAT also will be used on the 200 Mammals Project, which aims to add approximately 140 new mammalian genome assemblies to the existing set (https://karlssonlab.org/2017/08/03/the-200-mammals-project/). These projects will provide a new understanding of gene evolution.

## Materials and Methods

CAT produces as output a series of diagnostic plots, an annotation set for each target genome, and a UCSC comparative assembly hub^40^. Both the pipeline and associated documentation can be found at https://github.com/ComparativeGenomicsToolkit/Comparative-Annotation-Toolkit. CAT is constructed using the Luigi (https://github.com/spotify/luigi) workflow manager, with Toil^50^ used for computationally intensive steps that work best when submitted to a compute cluster.

### RNA-seq

CAT annotation is improved when species-specific RNA-seq data are provided. These data are used as hints for AugustusTMR and AugustusCGP. In AugustusTMR, RNA-seq helps fill in missing information in the alignment, as well as resolve evolutionary changes. In AugustusCGP, RNA-seq additionally helps prevent false positives inherent in *ab initio* gene finding. For these reasons, RNA-seq was obtained from SRA for all species annotated in this paper. All RNA-seq were aligned to their respective genomes with STAR^51^ and the resulting BAM files passed to CAT to construct the extrinsic hints database. See Table S2 for accessions and tissue types of RNA-seq data used for annotation. In addition, for the PacBio great ape annotation, RNA-seq data were generated using iPSC lines for human, chimpanzee, gorilla and orangutan derived from cells for the same individuals as the assemblies (Kronenberg et al., submitted). For all expression analyses, Kallisto^32^ was used.

### Annotation set similarity analysis

Jaccard similarity analysis was performed with BEDTools^52^. The rat locus overlap analysis was performed with the Kent tool clusterGenes, which requires exonic overlap on the same strand.

### Iso-Seq

Iso-Seq full-length non-chimeric reads (FLNC) were also generated from the great ape iPSC lines and aligned with GMAP^53^.To perform isoform-level validation in the primates, the Iso-Seq data used as input to CAT were also clustered with isoform-level clustering (ICE) and then collapsed into isoforms using ToFu^29^. Ensembl provided a pre-release of their new V91 annotations for panTro4 and gorGor4, but did not yet run their updated pipeline on ponAbe2.

### ICE validation

The output transcripts from ICE were compared to various annotation sets in both an exact and fuzzy matching scheme. In the exact scheme, the genomic order and positions of all of the introns (an intron chain) of a transcript are compared to any ICE isoforms which overlap it. In the fuzzy matching scheme, each annotated intron chain is allowed to move up to 8 bases in either direction and still be called a match.

### BUSCO

The mammalian BUSCO analysis was performed using the mammalia odb9 set of 4,104 genes. BUSCO was ran against the complete protein-coding sequence repertoire produced by CAT in that species in the ‘protein’ mode.

### Progressive Cactus

All Cactus alignments except the 14-way mammal alignment were generated using Progressive Cactus (https://github.com/glennhickey/ProgressiveCactus) commit 91d6344. For the mouse-rat alignment, the guide tree was

~~~
(((Lesser_Egyptian_jerboa:0.1,(Mouse:0.084509,Rat:0.091589)mouse_rat:0.107923)rodent
   :0.148738,Rabbit:0.21569)glires:0.015313,Human:0.143908);
~~~

For the primate alignments, the guide tree was

~~~
(((((((Susie_Gorilla:0.008964,(Human:0.00655,Clint_Chimp:0.00684)human_chimp:0.00122)
   gorilla_chimp_human:0.009693,Susie_Orangutan:0.01894)great_ape:0.003471,Gibbon:0.02227)
   great_ape_gibbon:0.01204,Rhesus:0.004991)old_world_monkey:0.02183,Squirrel_monkey
   :0.01035)monkey:0.05209,Bushbaby:0.1194)primate_anc:0.013494,Mouse:0.084509);
~~~

An identical tree (with different assembly names) was used for the alignment of current reference great apes.

For the diploid human alignments, the two haploid cell lines (PacBio) or all human haplotypes (10x) were placed under the same node with a very short branch length, with chimpanzee as outgroup. The guide trees were

~~~
((hg38:0.001,chm1:0.001,chm13:0.001)human:0.01,chimp:0.01);
~~~

and

~~~
((hg38:0.001,HG00512-H1:.001,HG00512-H2:.001,NA12878-H1:.001,NA12878-H2:.001,NA19240-H1:.001,
   NA19240-H2:.001,NA24385-H1:.001,NA24385-H2:.001)human:0.01,chimp:0.01);
~~~

representing a star phylogeny of the three human assemblies. For the 14-way mammal alignment, the Progressive Cactus commit used was e3c6055 and the guide tree was

~~~
((((oryCun2:0.21,((Pahari_EiJ:0.03,mm10:0.025107)1:0.02,rn6:0.013)1:0.252)1:0.01,((((hg19
   :0.00642915,panTro4:0.00638042)1:0.00217637,gorGor3:0.00882142)1:0.00935116,ponAbe2
   :0.0185056)1:0.00440069,rheMac3:0.007)1:0.1)1:0.02,((oviAri3:0.019,bosTau8:0.0506)
   1:0.17,(canFam3:0.11,felCat8:0.08)1:0.06)1:0.02)1:0.02,loxAfr3:0.15);
~~~

Slightly out-of-date versions of some assemblies (hg19 and rheMac3) were used because a collaborator had data on those assemblies that they wished to use the alignment to analyze. The rodent and primate subtrees were first aligned separately (the rodent subtree originally included additional mouse strains)^49, 54^. The two subtrees were then “stitched” together into a single alignment by aligning together their roots along with several Laurasiatheria genomes. This was done to save alignment time by reusing existing alignments.

### CAT

CAT was run on the UCSC Genome Browser compute cluster for all annotation efforts in this publication. CAT commit f89a814 was used. For a detailed description of how CAT works, see both the supplementary text as well as the README.md on github (https://github.com/ComparativeGenomicsToolkit/Comparative-Annotation-Toolkit).

### Pipeline runtime

CAT is relatively efficient, taking on the order of thousands of core hours to run. The largest considerations for runtime are running the various parameterizations of AUGUSTUS as well as generating the required cactus alignment. AugustusCGP may run significantly faster on alignments with many genomes by reducing the chunk size from the default, but at the cost of lower quality predictions. AugustusTMR runtime scales linearly with the number of protein-coding transcripts in the input annotation set, but scales non-linearly with the number of extrinsic hints provided, particularly if the hints are contradictory.

All of the analyses in this paper were run on the UCSC cluster, which uses the cluster management tool Parasol and has 1024 cores with 8GB of RAM per core. CAT was optimized for this, and shouldn’t need more memory per core in any case except the AugustusCGP step when the number of aligned genomes exceeds 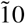. This can be adjusted by reducing the alignment chunk size that AugustusCGP is given to work with. For example, for the 14-way mammalian analysis, the flags `--maf-chunksize 1000000 --maf-overlap 200000` were set which kept memory usage under 8GB.

Cactus alignments take on the order of 120 CPU days (2,880 core hours) per internal node on the guide tree, assuming a binary tree. This number can fluctuate by a factor of 2-4 depending on how similar the two genomes being aligned at that node are. Cactus alignments are a mix of high CPU low memory steps with a few high memory steps, with some jobs requiring 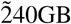 of RAM.

Running CAT on the PacBio primate genomes took a total of 7,030 core hours, with 3,437 of those dedicated to running AugustusTMR, 1,191 dedicated to running AugustusPB, and 2,190 dedicated to running AugustusCGP. Running CAT on the 14-way mammalian alignment took a total of 24,122 core hours, with 14,045 of those dedicated to running AugustusTMR, and 8,225 dedicated to running AugustusCGP.

## Acknowledgements

We would like to thank Brian Raney, Hiram Clawson and the rest of the UCSC Genome Browser team for help creating browser tracks. We would also like to thank James Kent for allowing UCSC Genome Browser compute resources to be used for this project. Finally, we would like to thank Fergal Martin and Paul Flicek for revising the paper and providing a pre-release of the Ensembl V91 annotations on chimpanzee and gorilla, as well as the whole GENCODE consortium for their support and advice. This work was supported, in part, by US National Institutes of Health (NIH) grants U24HG009081 and U41HG007635 to E.E.E., HG007990 to D.H and B.P, and HG007234 to B.P. E.E.E. and D.H. are investigators of the Howard Hughes Medical Institute. (COI: E.E.E. is on the scientific advisory board (SAB) of DNAnexus, Inc.)

